# Energetic determinants of the Par-3 interaction with the Par complex

**DOI:** 10.1101/2021.12.10.472131

**Authors:** Rhiannon R. Penkert, Elizabeth Vargas, Kenneth E. Prehoda

**Affiliations:** Institute of Molecular Biology, Department of Chemistry and Biochemistry, 1229 University of Oregon, Eugene, OR 97403

## Abstract

The animal cell polarity regulator Par-3 recruits the Par complex (Par-6 and atypical Protein Kinase C–aPKC) to specific sites on the cell membrane. Although numerous physical interactions have been reported between Par-3 and the Par complex, it has been unclear how each contributes to the overall interaction. Using purified, intact Par complex and a quantitative binding assay, we found that energy for this interaction is provided by Par-3’s second and third PDZ protein interaction domains. Both Par-3 PDZ domains bind to aPKC’s PDZ Binding Motif (PBM) in the Par complex, with binding energy contributed from aPKC’s adjacent catalytic domain. In addition to highlighting the role of Par-3 PDZ interactions with the aPKC kinase domain and PBM in stabilizing Par-3 – Par complex assembly, our results indicate that each Par-3 molecule can potentially recruit two Par complexes to the membrane during cell polarization.

## Introduction

The Par complex polarizes diverse animal cells by forming a specific domain on the plasma membrane. In the Par domain, the Par complex component atypical Protein Kinase C (aPKC) phosphorylates and displaces substrates, thereby restricting them to a complementary membrane domain (1). In this manner, the cellular pattern formed by Par-mediated polarity is ultimately determined by the mechanisms that target the Par complex to the membrane. Membrane recruitment relies at least in part on interactions with proteins that directly associate with the membrane, including the Rho GTPase Cdc42 and the multi-PDZ protein Par-3. The Par complex’s interaction with Cdc42 is via a single well-defined site, the Par complex component Par-6’s semi-CRIB domain (2–6). However, numerous interactions between Par-3 and the Par complex have been reported (2, 3, 7–10) and it has been unclear how each contributes to the overall interaction.

The interaction between Par-3 and the Par complex was originally discovered in the context of the interaction between aPKC and its phosphorylation site on Par-3, the aPKC Phosphorylation Motif (APM aka Conserved Region 3–CR3) (7). Subsequently, interactions were reported outside of aPKC’s catalytic domain: i) Par-3 PDZ1 and the Par-6 PDZ (2, 3, 11), ii) Par-3 PDZ2-3 acting together and aPKC (8), iii) Par-3 PDZ1 or PDZ3 binding to the Par-6 PDZ Binding Motif (PBM) (9), and iv) PDZ2 with a PBM in aPKC (10). Each of these interactions, except for the interaction of aPKC’s kinase domain with its substrate sequence on Par-3, involves one or more of Par-3’s three PDZ protein interaction domains.

Several factors have made it difficult to understand how the many interactions identified between Par-3 and the Par complex contribute to the overall interaction. Most interactions have not been examined in the context of the intact Par complex. In this context it is not possible to understand how individual interactions contribute to the overall energetics of Par-3 assembly with the Par complex, or if interactions might cooperate or compete. Furthermore, many of the interactions have not been examined quantitatively so it has not been possible to assess their relative strength. Finally, the presence of multiple potential Par complex binding sites on Par-3 raises the possibility that each Par-3 protein might bind more than one Par complex. Here we examine the energetics of Par-3 binding to the fully reconstituted Par complex using a quantitative binding assay to address these issues.

## Results

### Multiple interactions contribute to Par-3/Par complex interaction energy

To investigate the energetic determinants of Par-3’s interaction with the Par complex (Par-6 and aPKC), we measured binding energy using a supernatant depletion assay (Figure S1A). This assay uses solid (glutathione or amylose agarose resin) and soluble phases like a typical “pull-down” assay but the amount of protein in the soluble phase is monitored at equilibrium rather than what remains on the solid phase after washing (12). We used the PDZ1-APM region of Par-3 (Figure 1A) as a starting point because it contains all domains that have been reported to interact with the Par complex and it can be purified to a level suitable for quantitative analysis (all reagents used in this study are shown in Figure S1B) (10). We examined binding of Par-3 PDZ1-APM to the full Par complex to allow the multiple, potentially cooperative interactions to form. As shown in Figure 1B,C, the binding energy (ΔG°) of Par-3 PDZ1-APM to the Par complex is 8.6 kcal/mole (8.5-8.7 95% CI; all binding energies reported in this study can be found in Supplemental Table 1). Because of the potential for multiple interactions between Par-3 and the Par complex, this energy may be the cumulative effect of individual binding events. Below we examine how each of the potential interactions between Par-3 and the Par complex contributes to the overall binding energy.

**Figure 1.**
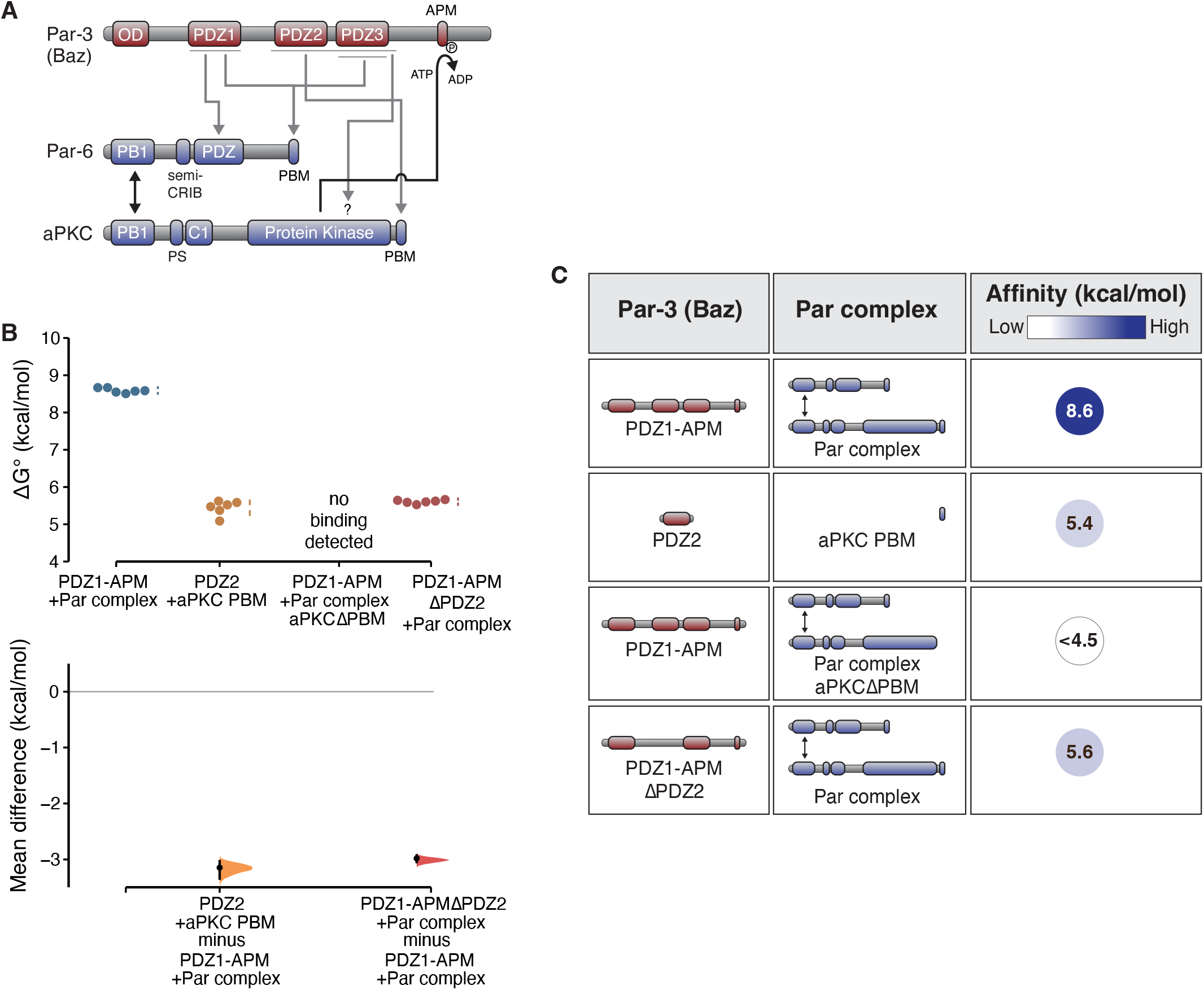
Energetic composition of the Par-3 interaction with the Par complex (**A**) Schematic of reported Par-3 interactions with the Par complex. (**B**) Cumming estimation plot of Par-3/Par complex interaction energies measured using the supernatant depletion assay. The result of each replicate is shown (filled circles) along with mean and standard deviation (gap and bars adjacent to filled circles) are shown in the top plot. The difference in the means relative to the PDZ1-APM/Par complex mean are shown in the bottom plot (filled circles) along with the 95% confidence intervals (black bar) derived from the bootstrap 95% confidence interval (shaded distribution). (**C**) Summary of binding energies for Par-3 and Par complex variants.

We recently discovered an interaction between the second of Par-3’s three PDZ domains (PDZ2) and a highly conserved PBM at the COOH-terminus of aPKC (10). The Par-3 PDZ2-aPKC PBM interaction is required for the recruitment of the Par complex to the cortex of asymmetrically dividing *Drosophila* neuroblasts. Using the supernatant depletion assay we found that this interaction has a binding energy of 5.4 kcal/mole (5.2-5.7 95% CI) which represents approximately 60% of the full Par-3 PDZ1-APM’s binding energy (Figure 1B,C). We were unable to detect an interaction between PDZ2 and a Par complex lacking aPKC’s PBM (the limit of detection of the supernatant depletion assay is approximately 4.5 kcal/mole) consistent with a central role of this motif in the overall interaction. Surprisingly, however, removal of PDZ2 did not abrogate binding as PDZ1-APM ΔPDZ2 bound the Par complex with approximately the same binding energy as that for Par-3 PDZ2–aPKC PBM (Figure 1B,C; 5.6 kcal/mole; 5.6-5.7 95% CI). We conclude that while Par-3 PDZ2–aPKC PBM represents a significant fraction of the Par-3 interaction with the Par complex, interactions outside of the PDZ2 (but also potentially involving the aPKC PBM) make a significant contribution. Furthermore, individual interactions appear to be non-additive (i.e. cooperative).

### The aPKC kinase domain and PBM form the Par complex binding surface for Par-3

We sought to determine which interaction domains or motifs from the Par complex collaborate with the aPKC PBM to contribute binding energy for Par-3. Par-6 has been reported to contain a PBM that interacts with Par-3 PDZ1 or PDZ3 (9). When examining the effect of removing Par-6’s PBM on the overall interaction energetics, we were unable to detect a difference in binding of Par-3 to the Par complex lacking the Par-6 PBM (Figure 2A,B; 8.6 kcal/mole; 8.5-8.8 95% CI). Given that the supernatant depletion assay reliably detects binding energy differences on the order of 0.2 kcal/mole, we conclude that Par-3 interactions with the Par-6 PBM do not play a significant role in stabilizing Par-3 binding to the Par complex in the context of these purified components.

**Figure 2.**
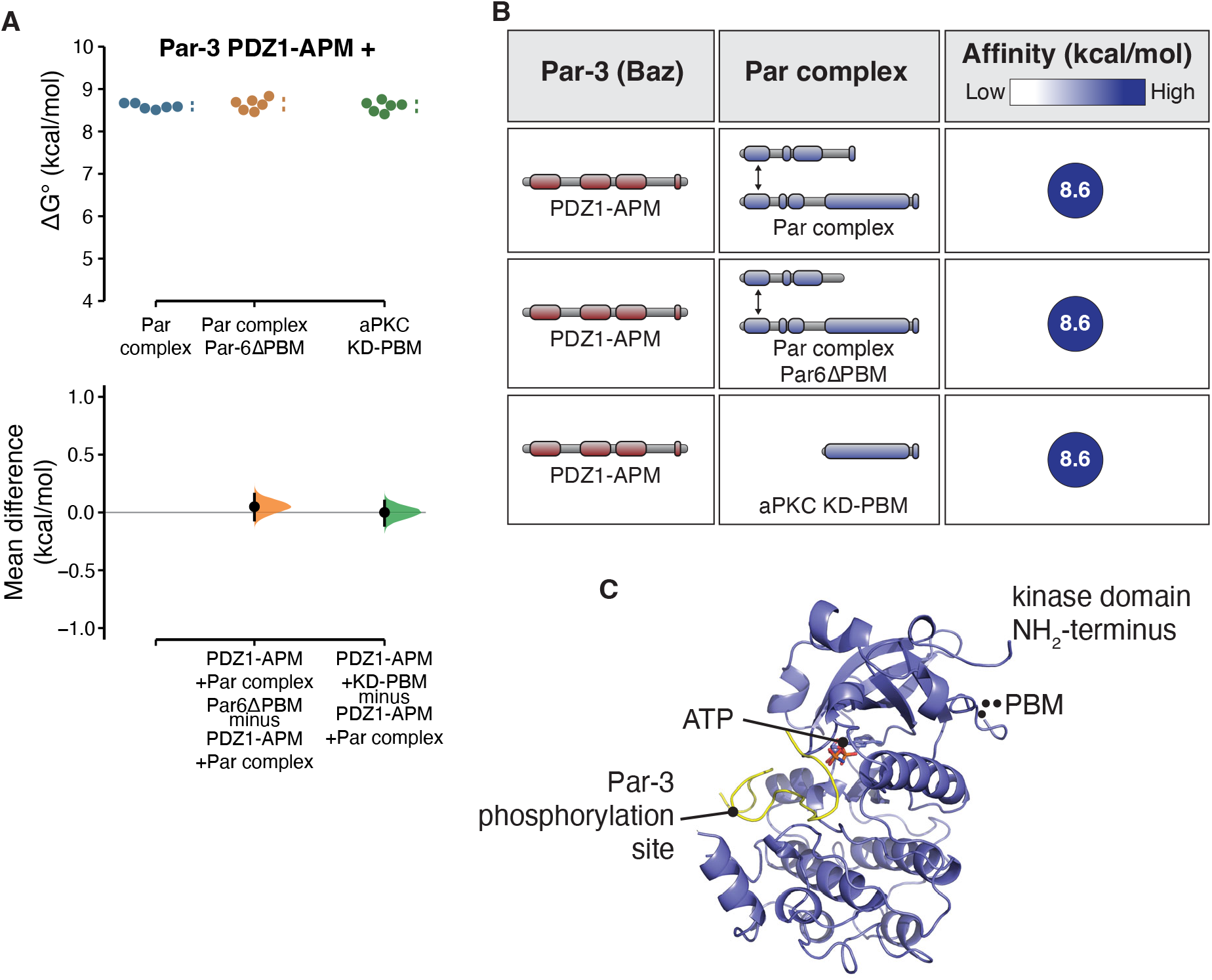
The aPKC kinase domain and PDZ binding motif form the Par complex binding site for Par-3 (**A**) Cumming estimation plot of Par-3/Par complex interaction energies measured using the supernatant depletion assay. Note: the data for PDZ1-APM binding to the Par complex is the same as shown in Figure 1. (**B**) Summary of binding energies for Par-3 interaction with the aPKC KD-PBM and Par complex lacking the Par-6 PBM. (**C**) Structure of the aPKC kinase domain in complex with the Par-3 phosphorylation site (from PDB ID 5LI1; Soriano et al 2016) showing the relative position of the PBM and substrate binding sites. Note that electron density for the residues directly preceding the PBM, but not the PBM itself, are present in this structure.

Given that the Par-6 PBM is not responsible for the additional interaction energy with Par-3, we sought to determine which Par complex interaction domains or motifs might contribute the additional binding energy beyond the aPKC PBM. We found that the aPKC kinase domain along with the adjacent PBM (KD-PBM; Figure 2C) fully recapitulated the interaction energy of the Par complex with Par-3 (Figure 2A,B; 8.6 kcal/mole; 8.5-8.7). Thus, in the context of these purified components, we do not find that the Par-3 PDZ1 interaction with the Par-6 PDZ, or the PDZ1 and 3 interactions with the Par-6 PBM substantially contribute to the overall Par-3 and Par complex binding energy.

### A conserved Basic Region NH_2_-terminal to Par-3 PDZ2 contributes to Par complex binding

We used both the Par complex and the isolated aPKC KD-PBM to identify which regions of Par-3 besides PDZ2 contribute to the overall interaction energy. We found that a Par-3 fragment containing its three PDZ domains has similar binding energy as PDZ1-APM (Figure 3A-D; 8.8 kcal/mole; 8.6-9.0 95% CI). This result indicates that Par-3’s phosphorylation site (APM) and the linker region connecting it to PDZ3 do not contribute significantly to the interaction. Note that ATP was present in our binding assay so that any interaction of the aPKC kinase domain with the APM was likely transient (and the interaction with the phosphorylated APM is weak) (10, 13).

**Figure 3.**
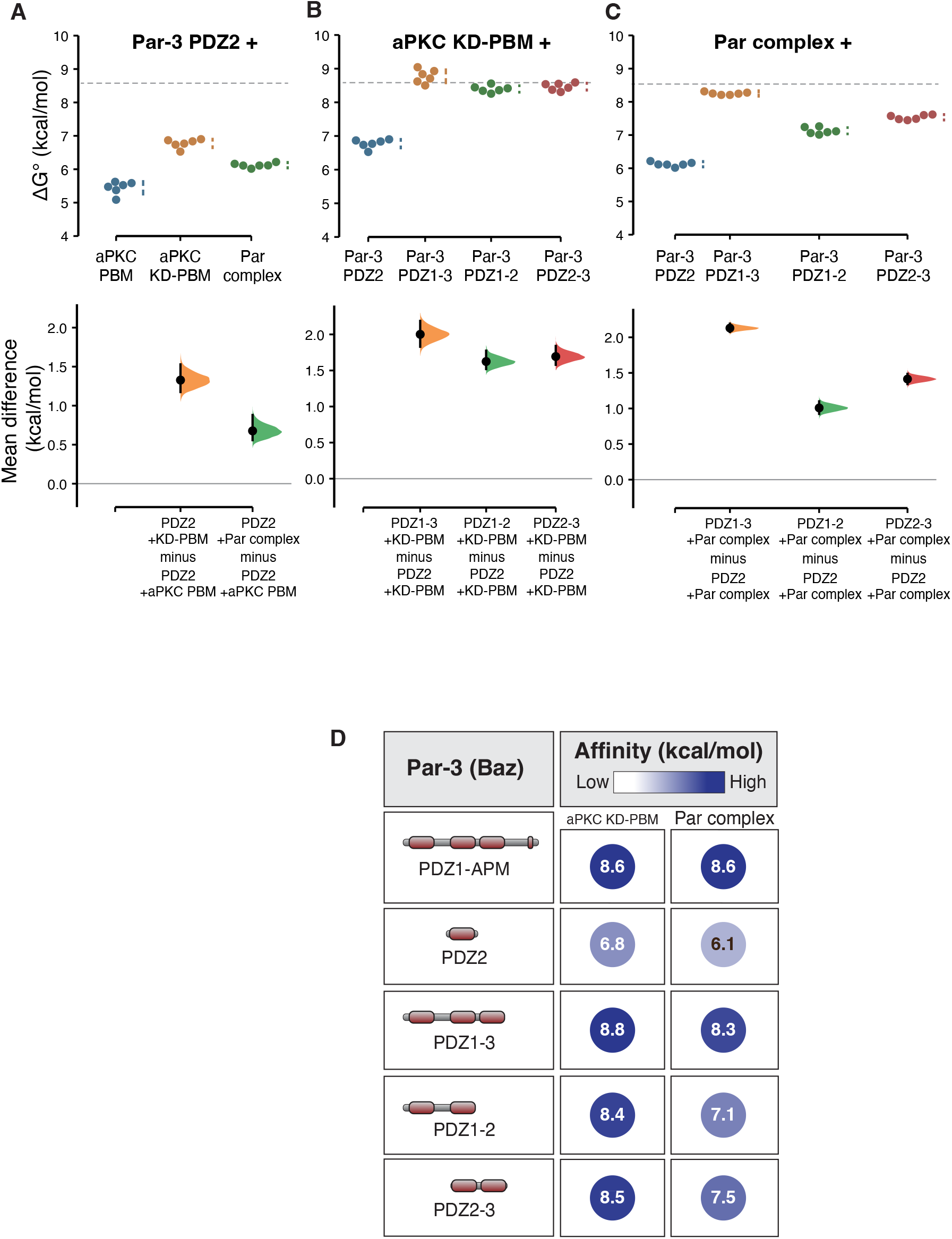
Energetic contributions to the Par-3/Par complex interaction from the Par-3 PDZ domains (**A-C**) Cumming estimation plots of Par-3/Par complex interaction energies measured using the supernatant depletion assay. The dashed lines indicate the binding energy of PDZ1-APM binding to the Par complex. (**D**) Summary of binding energies for Par-3 interaction with the aPKC KD-PBM and Par complex.

When examining the three Par-3 PDZ domains we found that either PDZ1-2 or PDZ2-3 bound with binding energies similar to PDZ1-APM although PDZ1-2’s was somewhat lower than PDZ2-3’s, an effect that was larger in the context of the full Par complex relative to the KD-PBM alone (Figure 3). We noticed that an approximately 30 residue sequence directly NH_2_-terminal to the PDZ2 domain is enriched in basic amino acids and highly conserved in Par-3’s from diverse animal species (Figure 4A). We termed this motif the Basic Region (BR) and found that including it with Par-3 PDZ2 (BR-PDZ2) significantly increased the binding energy of the interaction with the aPKC kinase domain and the full Par complex (Figure 4B,C). We conclude that the higher binding energy of PDZ1-2 compared to PDZ2 alone is contributed by the conserved BR motif.

**Figure 4.**
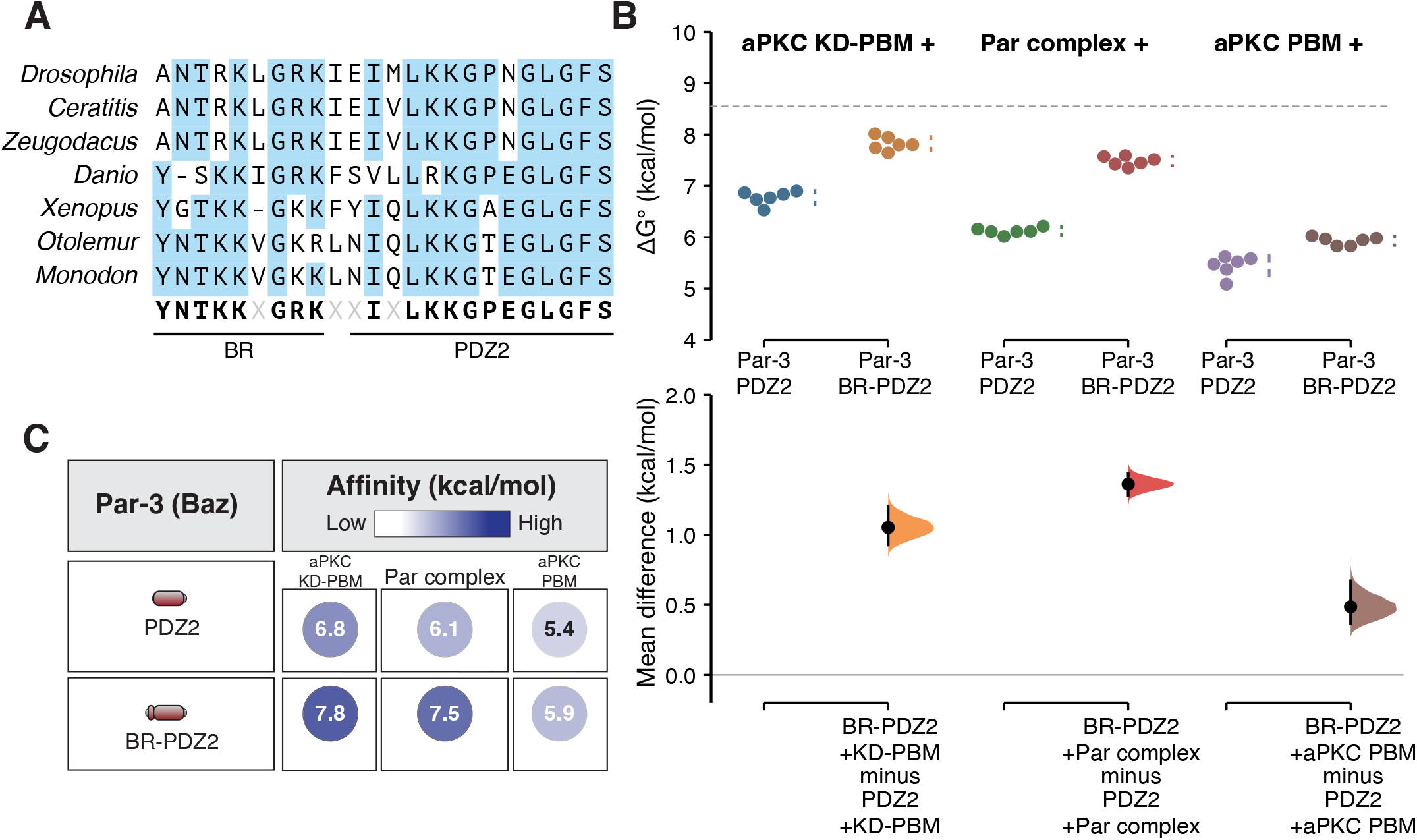
A conserved basic region (BR) contributes binding energy to the Par-3 PDZ2 interaction with the Par complex (**A**) Sequence alignment of the region NH_2_-terminal to the Par-3 PDZ2 from the Par-3 sequence from diverse animal species. (**B**) Cumming estimation plot of Par-3/Par complex interaction energies measured using the supernatant depletion assay. The dashed lines indicate the binding energy of PDZ1-APM binding to the Par complex. (**C**) Summary of binding energies for Par-3 PDZ2 and BR-PDZ2 interaction with the aPKC KD-PBM, Par complex, and aPKC PBM.

### The Par-3 PDZ3 domain binds the aPKC kinase domain and PBM

Like PDZ1-2, the combination of PDZ2 and 3 (Par-3 PDZ2-3) also bound aPKC KD-PBM and full Par complex with higher affinity than PDZ2 alone (Figure 5A-D). In this case, the higher binding energy originates from PDZ3 as we discovered that it binds the aPKC KD-PBM with similar energy to PDZ2 and somewhat less energy to the full Par complex (Figure 5A-D). We also found that PDZ3 binds the aPKC PBM with a similar energy as PDZ2. Like PDZ2, the binding energy of PDZ3 was higher for aPKC KD-PBM compared to the PBM alone.

**Figure 5.**
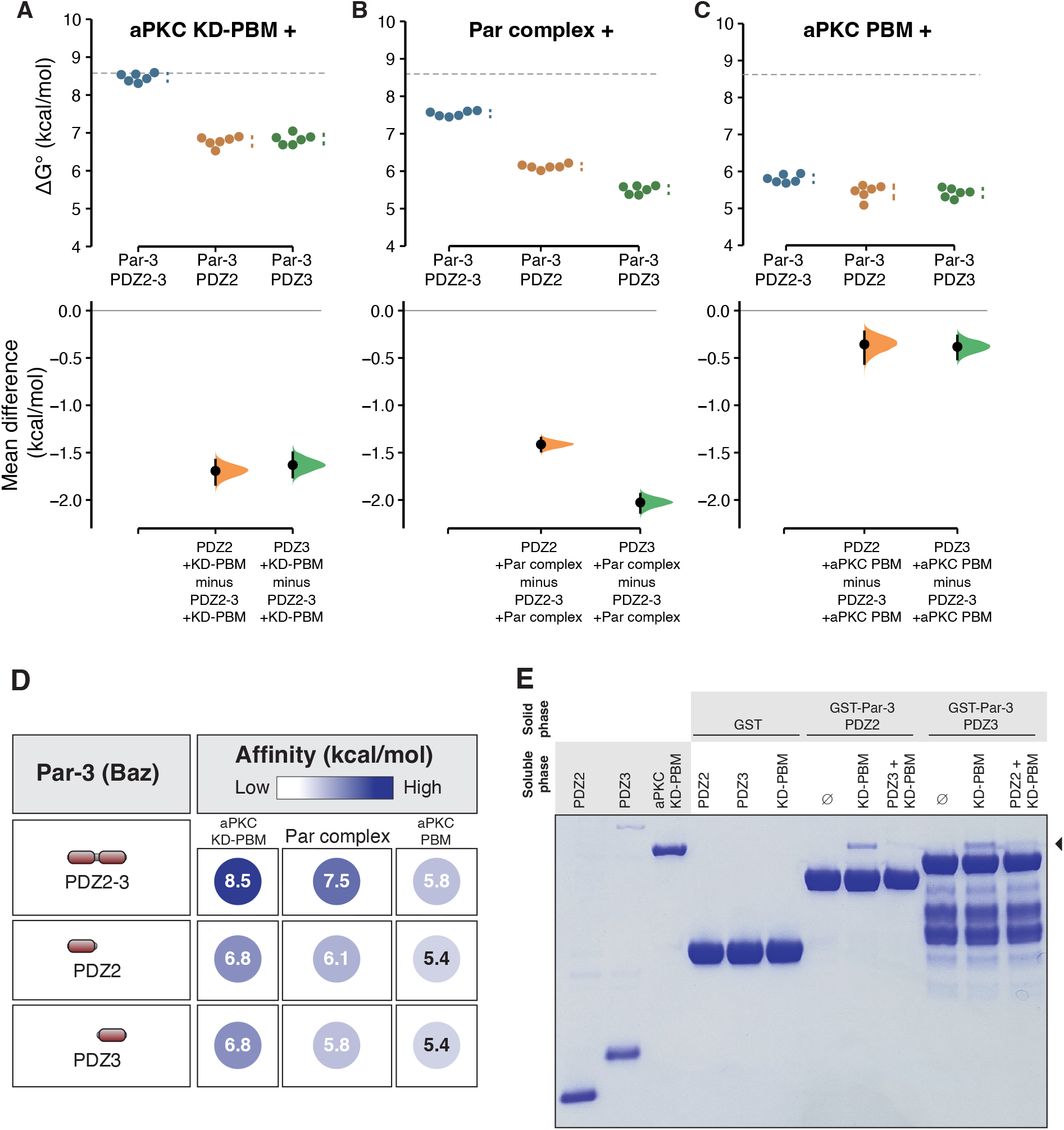
Par-3 PDZ3 binds the aPKC kinase domain and PDZ Binding Motif (**A-C**) Cumming estimation plots of Par-3 PDZ2 and PDZ3 interaction energies with the aPKC kinase domain–PBM (A), full Par complex (B) and aPKC PBM (C) measured using the supernatant depletion assay. Dashed lines represent the interaction energy of PDZ1-APM to the Par com- plex. (**D**) Summary of binding energies for Par-3 PDZ2 and PDZ3 interaction with the aPKC kinase domain-PBM, Par complex, and aPKC PBM. (**E**) Competition between Par-3 PDZ2 and PDZ3 for binding to aPKC KD-PBM. Solid phase (glutathione resin) bound Glutathione-S-Trans- ferase (GST) fused Par-3 PDZ2 or PDZ3 incubated with aPKC KD-PBM (arrowhead) and the indicated competing PDZ domain. Shaded regions of legend indicate the fraction applied to the gel (soluble-phase or solid-phase components after mixing with soluble-phase components and washing).

Our results indicate that Par-3 PDZ2 and PDZ3 use a similar binding mode and therefore may compete for binding to aPKC KD-PBM. To test this hypothesis, we performed a competition experiment, first assembling a complex of the aPKC KD-PBM with PDZ2 and then adding PDZ3. We found that the presence of PDZ3 caused a significant decrease in the amount of aPKC bound to PDZ2 (Figure 5E). Soluble PDZ2 was also able to displace PDZ3 from aPKC KD-PBM (Figure 5E). The competitive binding for the two PDZ domains suggests that PDZ2 and PDZ3 each binds a distinct Par complex. Furthermore, the increased binding affinity when both PDZ2 and 3 are present (e.g. PDZ2-3) relative to the individual domains likely arises from an avidity effect in which more than one Par complex is participating in the interaction.

### Par-3 BR-PDZ2-3 binding to aPKC KD-PBM recapitulates the overall interaction energy

Taken together, our results suggest that the binding energy of the Par-3 interaction with the Par complex arises from separate interactions of the BR-PDZ2 and PDZ3 with the aPKC KD-PBM. As shown in Figure 6A,B, Par-3 BR-PDZ2-3 nearly completely recapitulates the binding energy of PDZ1-APM. We conclude that distinct interactions of BR-PDZ2 and PDZ3 with aPKC KD-PBM form the basis of the Par-3 interaction with the Par complex (Figure 6C). In the context of these purified proteins, we do not detect a significant contribution from the interaction of PDZ1 or PDZ3 with the Par-6 PBM, the interaction of PDZ1 with the Par-6 PDZ, or the interaction of the aPKC kinase domain with its phosphorylation site.

**Figure 6.**
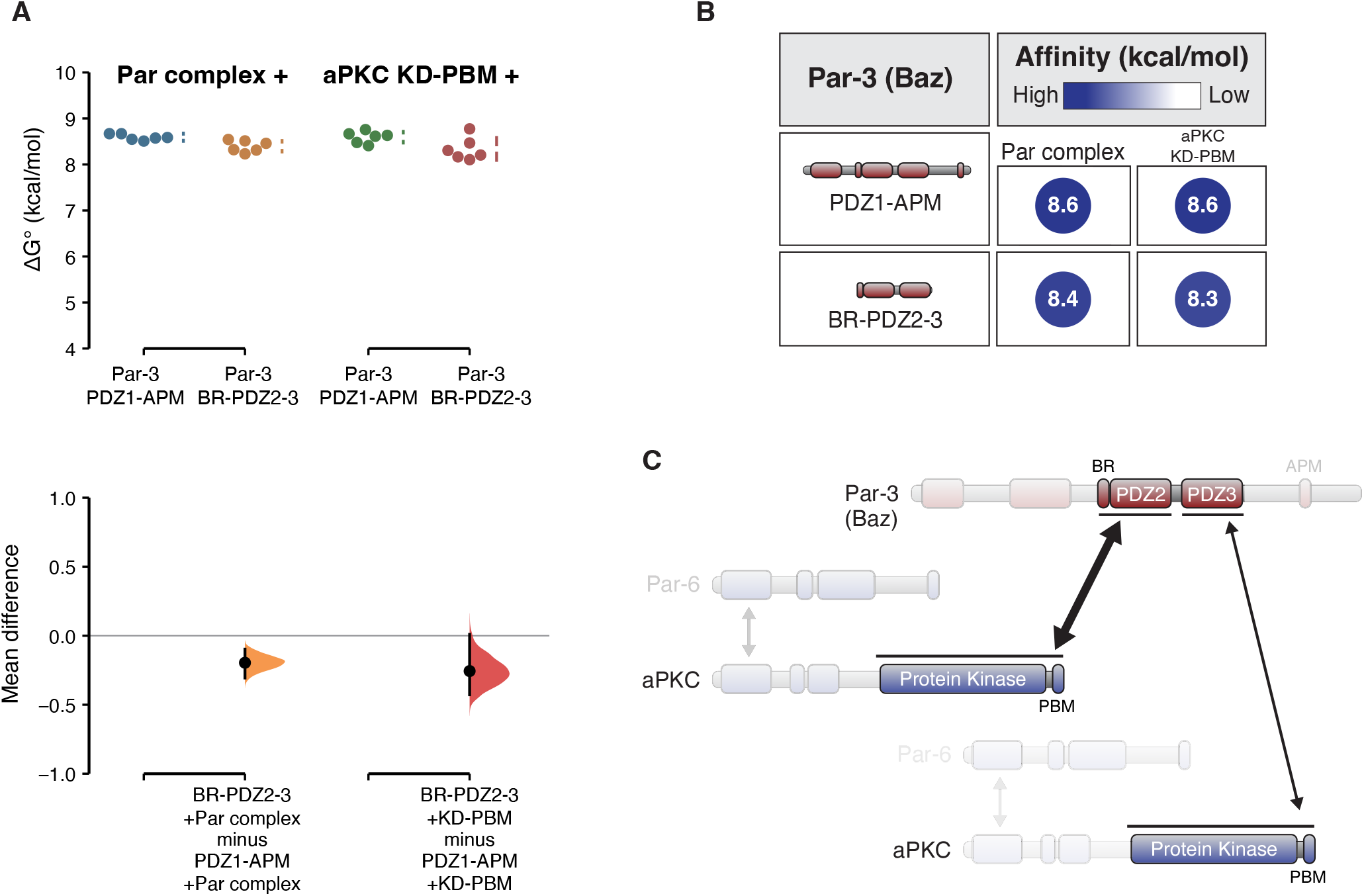
Par-3 BR-PDZ2-3 binding to the aPKC kinase domain-PBM fully recapitulates the Par-3 interaction with the Par complex (**A**) Cumming estimation plot of Par-3 BR-PDZ2-3 interaction energies with the full Par complex and aPKC KD-PBM measured using the supernatant depletion assay. (**B**) Summary of binding energies for BR-PDZ2-3 interaction with the full Par complex and aPKC KD-PBM. (**C**) Model for interaction of Par-3 with the Par complex.

## Discussion

The nature of the Par-3 interaction with the Par complex has been enigmatic (14–18). In this study, we used a quantitative biochemical approach with purified, full-length Par complex and a region of Par-3 that contains all known binding motifs to address the challenge of understanding this complicated interaction. We found that Par-3 PDZ2 and PDZ3 binding to the aPKC KD-PBM nearly fully recapitulates the binding energy of the overall interaction between Par-3 and the Par complex. We note that these interactions most closely resemble the previously identified interaction of Par-3 PDZ2-3 with full-length aPKC using a yeast two-hybrid assay (8). Here we examine the implications of our quantitative findings on Par-3’s role in Par-mediated polarity.

We used binding energy to evaluate the relative contribution of each of the identified Par-3 interactions with the Par complex. The binding energies of several of the interactions in the context of isolated Par complex fragments have been previously reported. The interaction of the aPKC kinase domain with the Par-3 APM has been reported to be very high (8.6 kcal/mole) (19). However, this interaction was measured in the absence of ATP, which prevents substrate turnover and is therefore not physiologically relevant (10, 13). We did not detect any contribution to the overall interaction between Par-3 and the Par complex from the Par-3 APM when ATP was present. The interactions of Par-3 PDZ1 and PDZ3 with the Par-6 PBM were measured using NMR and were found to be weak (5.0 and 5.8 kcal/mole, respectively). While these affinities are low, they are above the limit of detection of the supernatant depletion assay. However, we did not detect any significant contribution from the Par-6 PBM in the context of Par complex binding to Par-3; we did not detect an interaction of Par-3 with Par-6/aPKCΔPBM nor did we detect a change in affinity when the Par-6 PBM was removed (i.e. Par-6ΔPBM/aPKC).

An analysis of the Par-3 domains required for polarity in the *C. elegans* zygote found that PDZ1 and 3 were dispensable but the oligomerization domain and PDZ2 were necessary (20). A similar analysis found that the interaction of the Par-6 PDZ domain with Par-3 was also dispensable for Par complex function (11). An examination of the Par-6 PBM found that it is not required for viability in *Drosophila* and its removal did not have a measurable effect on Par-6 recruitment to the cortex of the embryonic epithelium except when the Par-6 PDZ was also removed (9). In a study of aPKC PBM function, aPKCΔPBM was not polarized to the apical membrane during the asymmetric division of *Drosophila* larval neuroblasts (10). These functional results are consistent with the primacy of the aPKC PBM in binding to Par-3. They also suggest that the biochemical redundancy between Par-3 PDZ2 and PDZ3 does not translate to *in vivo* function, either because of the lower affinity of PDZ3 or because PDZ2 participates in other essential functions besides binding to the Par complex.

Our results indicate that the aPKC kinase domain participates in the interaction with Par-3 PDZ2 and PDZ3. The nature of this interaction is not known but the proximity of the aPKC PBM to the kinase domain is suggestive (Figure 2C) (19). Binding to the PBM would bring PDZ surfaces outside of the PBM pocket near the kinase domain and could lead to so-called “docking” interactions that occur between protein kinase substrates and regions away for the kinase domain active site (21). Future efforts will be directed at exploring the nature of this interaction and any role it may have in regulating aPKC activity.

## Materials and Methods

### Cloning

GST-, MBP- and his-tagged Par-3 constructs, GST aPKC PBM and his aPKC kinase domain-PBM (residues 259-606) were cloned as previously described (10) using Gibson cloning (New England BioLabs), Q5 mutagenesis (New England BioLabs) or traditional methods. In addition to an N-terminal MBP tag, the Par-3 PDZ1-APM (residues 309-987) construct also contained a C-terminal his-tag. Par complex components (aPKC and his-Par-6) were cloned into pCMV as previously described (10, 22). Please see the Key Resources table for additional information on specific constructs.

### Expression

All proteins, except for Par complex constructs, were expressed in *E. coli* (strain BL21 DE3). Constructs were transformed into BL21 cells, grown overnight at 37°C on LB + ampicillin (Amp; 100 μg/mL). Resulting colonies were selected and used to inoculate 100mL LB+Amp starter cultures. Cultures were grown at 37°C to an OD_600_ of 0.6-1.0 and then diluted into 2L LB+Amp cultures. At an OD_600_ of 0.8-1.0 expression was induced with 0.5 mM IPTG for 2-3 hours. Cultures were centrifuged at 4,400g for 15 minutes to pellet cells. Media was removed and pellets were resuspended in nickel lysis buffer [50mM NaH_2_PO_4_, 300 mM NaCl, 10 mM Imidazole, pH 8.0], GST lysis buffer [1XPBS, 1 mM DTT, pH 7.5] or Maltose lysis buffer [20 mM Tris, 200 mM NaCl, 1 mM EDTA, 1 mM DTT, pH 7.5], as appropriate. Resuspended pellets were frozen in liquid N_2_ and stored at -80°C.

Par complex constructs were expressed in HEK 293F cells (Thermofisher), as previously described (10, 22). Briefly, cells were grown in FreeStyle 293 expression media (Thermofisher) in shaker flasks at 37°C with 8% CO_2_. Cells were transfected with 293fectin (Thermofisher) or ExpiFectamine (Thermofisher) according to the manufacture’s protocol. After 48 hours, cells were collected by centrifugation (500g for 5 min). Cell pellets were resuspended in nickel lysis buffer, frozen in liquid N_2_ and stored at -80°C.

### Purification

Resuspended *E*.*coli* pellets were thawed and cells were lysed by probe sonication using a Sonicator Dismembrator (Model 500, Fisher Scientific; 70% amplitude, 0.3/0.7s on/off pulse, 3×1 min). 293F cell pellets were lysed similarly using a microtip probe (70% amplitude, 0.3/0.7s on/off pulse, 4×1 min). Lysates were centrifuged at 27,000g for 20 min to pellet cellular debris. GST- and MBP-tagged protein lysates were aliquoted, frozen in liquid N_2_ and stored at -80°C.

His-tagged protein lysates, except for aPKC KD-PBM, were incubated with HisPur Ni-NTA (Thermofisher) or HisPur Cobalt (Thermofisher) resin for 30 min at 4°C and then washed 3x with nickel lysis buffer. For 293F lysates, 100μM ATP and 5mM MgCl_2_ were added to the first and second washes. Proteins were eluted in 0.5-1.5mL fractions with nickel elution buffer (50 mM NaH_2_PO_4_, 300 mM NaCl, 300 mM Imidazole, pH 8.0). For all proteins, aside from Par complex, fractions containing protein were pooled, buffered exchanged into 20mM HEPES pH 7.5, 100 mM NaCl and 1 mM DTT using a PD10 desalting column (Cytiva), concentrated using a Vivaspin20 protein concentrator spin column (Cytiva), aliquoted, frozen in liquid N_2_ and stored at -80°C. For Par complex, proteins were further purified using anion exchange chromatography on an AKTA FPLC protein purification system (Amersham Biosciences). Following his-purification fractions were pooled and buffered exchanged into 20mM HEPES pH 7.5, 100 mM NaCl, 1 mM DTT, 100 μM ATP and 5 mM MgCl_2_ using a PD10 desalting column (Cytiva). Buffer-shifted protein was injected onto a Source Q (Cytiva) column and eluted over a salt gradient of 100-550mM NaCl. Fractions containing Par complex were pooled, buffered exchanged into 20 mM HEPES pH 7.5, 100 mM NaCl, 1 mM DTT, 100 μM ATP, and 5 mM MgCl_2_ using a PD10 desalting column (Cytiva), concentrated using a Vivaspin20 protein concentrator spin column (Cytiva), aliquoted, frozen in liquid N_2_ and stored at -80°C.

Due to solubility issues, aPKC KD-PBM was expressed in *E. coli* and his-purified partially under denaturing conditions. Following sonication and centrifugation (described above), the soluble fraction was discarded and the insoluble pellet was resuspended in 50mM NaH_2_PO_4_, 300 mM NaCl, 10 mM Imidazole, 8M Urea pH 8.0. Centrifugation was repeated (27,000g for 20 min) and the resulting soluble phase was incubated with HisPur Ni-NTA resin (ThermoFisher) for 30 min at 4°C. Resin was washed and eluted as described above. Purified protein was aliquoted, frozen in liquid N_2_ and stored at -80°C.

### Quantitative binding assay

For all solid phase proteins, except Par-3 PDZ1-APM and PDZ1-APMΔPDZ2, GST lysates were incubated with glutathione agarose resin (GoldBio; 50 μL resin per 0.5-1.5 mL of lysate) for 30 min at 4°C and then washed 6x (3x quick washes, followed by 3x 5min washes at room temp) with binding buffer (10 mM HEPES pH 7.5, 100 mM NaCl, 1 mM DTT 200 μM ATP, 5 mM MgCl_2_ and 0.1% Tween-20). After washing, resin was resuspended in 50 μL binding buffer to create a 50% slurry. Par-3 PDZ1-APM and PDZ1-APMΔPDZ2 were double tagged (N-terminal MBP-tag and C-terminal his-tag). Par-3 PDZ1-APM and PDZ1-APMΔPDZ2 were first his-purified (described above) prior to incubation with amylose resin (NEB). Amylose-bound Par-3 was then washed and resuspended in binding buffer as described for GST proteins.

Separately, unlabeled resin (amylose or glutathione resin, as appropriate) was washed 3x and resuspended in a 50% slurry with binding buffer. GST- or MBP-labeled resin was then serially diluted 1:1 (30 μL of 50% slurry) with unlabeled resin to create a gradient of the GST/MBP-tagged protein. Unlabeled resin was used as a negative control for binding. Soluble protein was added to the solid phase protein and incubated for one hour at room temperature with rotational mixing (see Supplemental Table 1 for the solid phase-soluble phase combination for each experiment), except for MBP-Par-3 constructs. Due to high levels of leaching into the supernatant from amylose bound MBP-tagged proteins, MBP-Par-3 assays were incubated for ten minutes (we confirmed that GST-Par-3 PDZ1-3 incubated for one hour produced indistinguishable results to MBP-Par-3 PDZ1-3 incubated for ten minutes).

Following incubation, a sample of the supernatant was removed from each tube and combined with 4X LDS sample buffer (ThermoFisher). Samples were run on a Bis-Tris gel, stained with Coomassie Brilliant Blue R-250 (GolBio) and band intensity was quantified using ImageJ (v1.53a). The fraction bound was determined using the following equation:

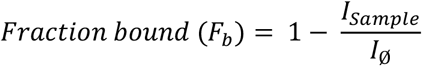

where I_sample_ is the intensity of the band at the current ligand concentration and I_ø_ is the band intensity without any ligand present. The dilution of the solid phase that resulted in 30-60% depletion (F_b_ = 0.3-0.6) was determined and the assay was repeated in sextuplicate at this dilution. The solid phase concentration ([L]) was determined by gel analysis using a standard protein of known concentration. For each repeat, the binding energy was then determined using the following the equation:

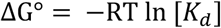

The binding dissociation constant (K_d_) was calculated using the following equation, assuming excess ligand conditions:

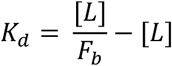

The data was visualized and analyzed using Excel (v16.53), GraphPad Prism (v9.2) and estimationstats.com.

### Qualitative binding assays

For GST pulldown assays, GST lysates were incubated with glutathione agarose resin (GoldBio) for 30 min at 4°C and then washed 6x (3x quick washes, followed by 3x 5 min washes at room temp) with binding buffer (10 mM HEPES pH 7.5, 100 mM NaCl, 1 mM DTT 200 μM ATP, 5 mM MgCl_2_ and 0.1% Tween-20). Soluble proteins were added to GST-bound proteins, as indicated, and incubated at room temperature with rotational agitation for 30-60 min. Resin was then washed 3x with binding buffer and protein was eluted with 4X LDS sample buffer (ThermoFisher). Samples were run on a Bis-Tris gel and stained with Coomassie Brilliant Blue R-250 (GolBio).

## Key Resources Table

| Reagent type (species) or resource | Designation | Source or reference | Identifiers | Additional information |
| --- | --- | --- | --- | --- |
| Recombinant protein | Par complex (his-Par-6, aPKC) | PMID: 32084408 |  | expressed in 293F cells from pCMV his-Par-6 1-351 and pCMV aPKC 1-606 |
| Recombinant protein | Par complex aPKCΔPBM | PMID: 32084408 |  | expressed in 293F cells from pCMV his-Par-6 1-351 and pCMV aPKC 1-600 |
| Recombinant protein | Par complex Par-6ΔPBM | PMID: 32084408 |  | expressed in 293F cells from pCMV his-Par-6 1-343 and pCMV aPKC 1-606 |
| Recombinant protein | GST-aPKC PBM | PMID: 32084408 |  | expressed in BL21 cells from pGEX aPKC 583-606 |
| Recombinant protein | GST-Par-3 PDZ2-3 | This paper |  | Cloned by Q5 mutagenesis; expressed in BL21 cells from pGEX Par-3 444-741 |
| Recombinant protein | GST-Par-3 PDZ1-2 | This paper |  | Gibson cloning; expressed in BL21 cells from pGEX Par-3 309-533 |
| Recombinant protein | GST-Par-3 PDZ1-3 | This paper |  | Gibson cloning; expressed in BL21 cells from pGEX Par-3 309-741 |
| Recombinant protein | GST-Par-3 BR-PDZ2 | This paper |  | Gibson cloning; expressed in BL21 cells from pGEX Par-3 426-533 |
| Recombinant protein | GST-Par-3 PDZ2 | This paper |  | Gibson cloning; expressed in BL21 cells from pGEX Par-3 444-533 |
| Recombinant protein | GST-Par-3 BR-PDZ2-3 | This paper |  | Gibson cloning; expressed in BL21 cells from pGEX Par-3 426-741 |
| Recombinant protein | GST-Par-3 PDZ3 | This paper |  | Gibson cloning; expressed in BL21 cells from pGEX Par-3 616-741 |
| Recombinant protein | aPKC KD-PBM | This paper |  | Cloned using traditional methods; expressed in BL21 cells from pBH aPKC |
|  |  |  |  | 259-606; his-purified under denaturing conditions |
| Recombinant protein | Par-3 PDZ2 | This paper |  | Gibson cloning; expressed in BL21 cells from pET19 Par-3 444-533; his-purified |
| Recombinant protein | Par-3 BR-PDZ2 | This paper |  | Gibson cloning; expressed in BL21 cells from pET19 Par-3 426-533; his-purified |
| Recombinant protein | Par-3 PDZ2-3 | This paper |  | Cloned by Q5 mutagenesis; expressed in BL21 cells from pET19 Par-3 444-741; his-purified |
| Recombinant protein | Par-3 PDZ3 | This paper |  | Gibson cloning; expressed in BL21 cells from pET19 Par-3 616-741; his-purified |
| Recombinant protein | MBP-Par-3 PDZ1-APM | PMID: 32084408 |  | expressed in BL21 cells from pMAL Par-3 309-987-his; C-terminal his-tag; his-purified prior to use in binding assay |
| Recombinant protein | MBP-Par-3 PDZ1-APMΔPDZ2 | PMID: 32084408 | | expressed in BL21 cells from pMAL Par-3 309-987Δ437-533-his; C-terminal his-tag; his-purified prior to use in binding assay |
| Recombinant DNA reagent | pCMV (mammalian expression plasmid) | ThermoFisher | 10586014 |  |
| Recombinant DNA reagent | pMal C4X (bacterial expression plasmid) | Addgene | 75288 |  |
| Recombinant DNA reagent | pGEX 4Ti (bacterial expression plasmid) | Amersham | 27458001 |  |
| Recombinant DNA reagent | pBH (bacterial expression plasmid) | PMID: 15023337 |  |  |
| Recombinant DNA reagent | pET19 (bacterial expression plasmid) | Millipore Sigma (Novagen) | 69677 |  |
| Bacterial strain | BL21-DE3 |  |  | used for recombinant protein expression |
| Bacterial strain | TG1 |  |  | used for DNA cloning |
| Cell line (human) | FreeStyle 293-F | ThermoFisher | R79007 |  |
| Chemical | IPTG | GoldBio | I2481C100 | used at 0.5mM |
| Chemical | 293fectin | ThermoFisher | 12347019 |  |
| Chemical | ExpiFectamine 293 Transfection Kit | ThermoFisher | A14524 |  |
| Chemical | Freestyle 293 expression Medium | ThermoFisher | 12338018 |  |
| Chemical | Opti-Mem | ThermoFisher | 3198588 |  |
| Chemical | HisPur cobalt resin | ThermoFisher | 89965 |  |
| Chemical | HisPur NiNTA resin | ThermoFisher | 88222 |  |
| Chemical | Amylose Resin | NEB | E8021L |  |
| Chemical | Glutathione Resin | GoldBio | G250-100 |  |
| Chemical | Source Q anion exchange resin | GE Healthcare | 17-1275-01 |  |
| Chemical | LB Broth, Miller | Millipore Sigma | 71753-6 |  |
| Chemical | 4X BOLT LDS sample buffer | ThermoFisher | B0007 |  |
| Chemical | 20X BOLT MES SDS running buffer | ThermoFisher | B0002 |  |
| Chemical | Coomassie Brilliant Blue R-250 | GoldBio | C-461-5 |  |
| Commercial kit | Q5® Site-Directed Mutagenesis Kit | NEB | E0552S |  |
| Commercial kit | Gibson Assembly Cloning Kit | NEB | E5510S |  |
| Other | Bolt 12% Bis-Tris Gels | ThermoFisher | NW00125BOX |  |
| Other | PD10 Desalting columns |  | 95017-001 |  |
| Other | VivaSpin 20 sample concentrators MWCO 30kD | Cytiva | 28932361 |  |
| Other | VivaSpin 20 sample concentrators MWCO 10kD | Cytiva | 28932360 |  |
| Other | VivaSpin 20 sample concentrators MWCO 5kD | Cytiva | 28932359 |  |
| Other | Shaker Flasks – 125mL | VWR | 89095-258 |  |
| Other | Shaker Flasks – 250mL | VWR | 89095-266 |  |
| software | ImageJ | NIH |  | v1.53a;<br>https://imagej.nih.gov/ij/ |
| software | GraphPad Prism | GraphPad Software |  | v.9.2 |
| software | Estimation Statistics BETA | PMID: 31217592 |  | www.estimationstats.com |

## Acknowledgments

This work was supported by NIH grant GM127092 (K.E.P.).

---

**Figure S1.**
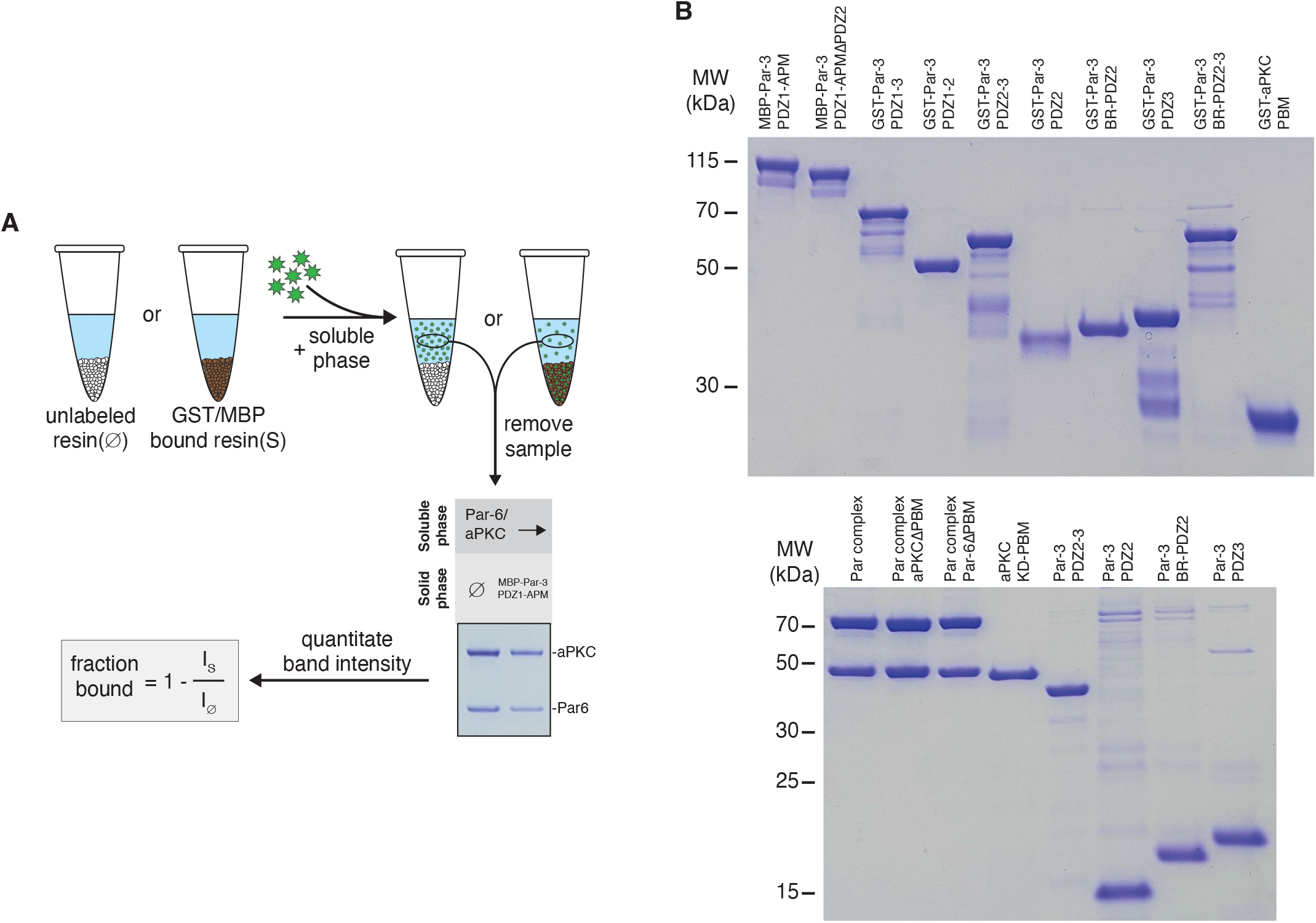
Supernatant depletion assay and reagents used (**A**) Schematic of the supernatant depletion quantitative binding assay. (**B**) Protein reagents used in this study

## References

1. Bailey, M. J., and Prehoda, K. E. (2015) Establishment of Par-Polarized Cortical Domains via Phosphoregulated Membrane Motifs. Dev. Cell. 35, 199–210

2. Joberty, G., Petersen, C., Gao, L., and Macara, I. G. (2000) The cell-polarity protein Par6 links Par3 and atypical protein kinase C to Cdc42. Nat. Cell Biol. 2, 531–539

3. Lin, D., Edwards, A. S., Fawcett, J. P., Mbamalu, G., Scott, J. D., and Pawson, T. (2000) A mammalian PAR-3-PAR-6 complex implicated in Cdc42/Rac1 and aPKC signalling and cell polarity. Nat. Cell Biol. 2, 540–547

4. Qiu, R. G., Abo, A., and Steven Martin, G. (2000) A human homolog of the C. elegans polarity determinant Par-6 links Rac and Cdc42 to PKCzeta signaling and cell transformation. Curr. Biol. CB. 10, 697–707

5. Noda, Y., Takeya, R., Ohno, S., Naito, S., Ito, T., and Sumimoto, H. (2001) Human homologues of the Caenorhabditis elegans cell polarity protein PAR6 as an adaptor that links the small GTPases Rac and Cdc42 to atypical protein kinase C. Genes Cells Devoted Mol. Cell. Mech. 6, 107–119

6. Garrard, S. M., Capaldo, C. T., Gao, L., Rosen, M. K., Macara, I. G., and Tomchick, D. R. (2003) Structure of Cdc42 in a complex with the GTPase-binding domain of the cell polarity protein, Par6. 22, 1125–1133

7. Izumi, Y., Hirose, T., Tamai, Y., Hirai, S., Nagashima, Y., Fujimoto, T., Tabuse, Y., Kemphues, K. J., and Ohno, S. (1998) An atypical PKC directly associates and colocalizes at the epithelial tight junction with ASIP, a mammalian homologue of Caenorhabditis elegans polarity protein PAR-3. J. Cell Biol. 143, 95–106

8. Wodarz, A., Ramrath, A., Grimm, A., and Knust, E. (2000) Drosophila atypical protein kinase C associates with Bazooka and controls polarity of epithelia and neuroblasts. J. Cell Biol. 150, 1361–1374

9. Renschler, F. A., Bruekner, S. R., Salomon, P. L., Mukherjee, A., Kullmann, L., Schütz-Stoffregen, M. C., Henzler, C., Pawson, T., Krahn, M. P., and Wiesner, S. (2018) Structural basis for the interaction between the cell polarity proteins Par3 and Par6. Sci. Signal. 10.1126/scisignal.aam9899

10. Holly, R. W., Jones, K., and Prehoda, K. E. (2020) A Conserved PDZ-Binding Motif in aPKC Interacts with Par-3 and Mediates Cortical Polarity. Curr. Biol. CB. 30, 893-898.e5

11. Li, J., Kim, H., Aceto, D. G., Hung, J., Aono, S., and Kemphues, K. J. (2010) Binding to PKC-3, but not to PAR-3 or to a conventional PDZ domain ligand, is required for PAR-6 function in C. elegans. Dev. Biol. 340, 88–98

12. Pollard, T. D. (2010) A guide to simple and informative binding assays. Mol. Biol. Cell. 21, 4061–4067

13. Holly, R. W., and Prehoda, K. E. (2019) Phosphorylation of Par-3 by Atypical Protein Kinase C and Competition between Its Substrates. Dev. Cell. 49, 678–679

14. Lang, C. F., and Munro, E. (2017) The PAR proteins: from molecular circuits to dynamic self-stabilizing cell polarity. Dev. Camb. Engl. 144, 3405–3416

15. Wu, X., Cai, Q., Feng, Z., and Zhang, M. (2020) Liquid-Liquid Phase Separation in Neuronal Development and Synaptic Signaling. Dev. Cell. 55, 18–29

16. Riga, A., Castiglioni, V. G., and Boxem, M. (2020) New insights into apical-basal polarization in epithelia. Curr. Opin. Cell Biol. 62, 1–8

17. Thompson, B. J. (2021) Par-3 family proteins in cell polarity & adhesion. FEBS J. 10.1111/febs.15754

18. Martin, E., Girardello, R., Dittmar, G., and Ludwig, A. (2021) New insights into the organization and regulation of the apical polarity network in mammalian epithelial cells. FEBS J. 10.1111/febs.15710

19. Soriano, E. V., Ivanova, M. E., Fletcher, G., Riou, P., Knowles, P. P., Barnouin, K., Purkiss, A., Kostelecky, B., Saiu, P., Linch, M., Elbediwy, A., Kjær, S., O’Reilly, N., Snijders, A. P., Parker, P. J., Thompson, B. J., and McDonald, N. Q. (2016) aPKC Inhibition by Par3 CR3 Flanking Regions Controls Substrate Access and Underpins Apical-Junctional Polarization. Dev. Cell. 38, 384–398

20. Li, B., Kim, H., Beers, M., and Kemphues, K. (2010) Different domains of C. elegans PAR-3 are required at different times in development. Dev. Biol. 344, 745–757

21. Reményi, A., Good, M. C., and Lim, W. A. (2006) Docking interactions in protein kinase and phosphatase networks. Curr. Opin. Struct. Biol. 16, 676–685

22. Graybill, C., Wee, B., Atwood, S. X., and Prehoda, K. E. (2012) Partitioning-defective protein 6 (Par-6) activates atypical protein kinase C (aPKC) by pseudosubstrate displacement. J. Biol. Chem. 287, 21003–21011

