## Supplemental Table 1 for "Energetic determinants of the Par-3 interaction with the Par complex"

|  | **Solid phase** | **Soluble phase** | **Mean  [95% CI]** |
| --- | --- | --- | --- |
| **PDZ1-APM +Par complex** | MBP-Par-3  309-987-his | aPKC 1-606  his-Par6 1-351 | 8.59  [8.52-8.67] |
| **PDZ2 +aPKC PBM** | GST-aPKC  583-606 | his-Par-3  444-533 | 5.44  [5.24-5.65] |
| **PDZ1-APM +Par complex aPKCΔPBM** | MBP-Par-3  309-987-his | aPKC 1-600  his-Par6 1-351 | No binding detected |
| **PDZ1-APM ΔPDZ2 +Par complex** | MBP-Par-3  309-987  Δ437-533 | aPKC 1-606  his-Par6 1-351 | 5.61  [5.56-5.66] |
| **PDZ1-APM +Par complex Par6ΔPBM** | MBP-Par-3  309-987-his | aPKC 1-606  his-Par6 1-343 | 8.64  [8.50-8.79] |
| **PDZ1-APM +KD-PBM** | MBP-Par-3  309-987-his | his-aPKC  259-606 | 8.59  [8.46-8.73] |
| **PDZ2 +KD-PBM** | GST-Par-3  444-533 | his-aPKC  259-606 | 6.77  [6.63-6.91] |
| **PDZ2 +Par complex** | GST-Par-3  444-533 | aPKC 1-606  his-Par6 1-351 | 6.12  [6.05-6.19] |
| **PDZ1-3 +KD-PBM** | GST-Par-3  309-741 | his-aPKC  259-606 | 8.77  [8.56-8.98] |
| **PDZ1-2 +KD-PBM** | GST-Par-3  309-533 | his-aPKC  259-606 | 8.40  [8.29-8.50] |
| **PDZ2-3 +KD-PBM** | GST-Par-3  444-741 | his-aPKC  259-606 | 8.47  [8.35- 8.58] |
| **PDZ1-3 +Par complex** | GST-Par-3  309-741 | aPKC 1-606  his-Par6 1-351 | 8.25  [8.21-8.30] |
| **PDZ1-2 +Par complex** | GST-Par-3  309-533 | aPKC 1-606  his-Par6 1-351 | 7.13  [7.02-7.24] |
| **PDZ2-3 +Par complex** | GST-Par-3  444-741 | aPKC 1-606  his-Par6 1-351 | 7.54  [7.46-7.61] |
| **BR-PDZ2 +KD-PBM** | GST-Par-3  426-533 | his-aPKC  259-606 | 7.82  [7.68-7.97] |
| **BR-PDZ2 +Par complex** | GST-Par-3  426-533 | aPKC 1-606  his-Par6 1-351 | 7.48  [7.39-7.58] |
| **BR-PDZ2 +aPKC PBM** | GST-aPKC  583-606 | his-Par-3  426-533 | 5.93  [5.84-6.02] |
| **PDZ3 +KD-PBM** | GST-Par-3  616-741 | his-aPKC  259-606 | 6.84  [6.69-6.98] |
| **PDZ3 +Par complex** | GST-Par-3  616-741 | aPKC 1-606  his-Par6 1-351 | 5.51  [5.39-5.63] |
| **PDZ3 +aPKC PBM** | GST-aPKC  583-606 | his-Par-3  616-741 | 5.42  [5.28-5.56] |
| **PDZ2-3 +aPKC PBM** | GST-aPKC  583-606 | his-Par-3  444-741 | 5.80  [5.69-5.92] |
| **BR-PDZ2-3 +KD-PBM** | GST-Par-3  426-741 | his-aPKC  259-606 | 8.34  [8.08-8.60] |
| **BR-PDZ2-3 +Par complex** | GST-Par-3  426-741 | aPKC 1-606  his-Par6 1-351 | 8.40  [8.26-8.53] |
